# A Deep Learning Framework for Predicting Gut Microbe-Host Receptor Interactions

**DOI:** 10.64898/2026.03.19.713066

**Authors:** Haimeng Li, Rui Zhao, Congmin Zhu, Rui Jiang, Ting Chen, Xiangyu Li, Yuqing Yang

## Abstract

**Motivation:** Gut microbiota regulates host health through complex protein-protein interactions. However, deciphering this specific interactions between microbiota and human receptors remains a significant challenge due to the lack of specialized computational tools.

**Results:** Leveraging the hypothesis of cell communication and relevant data, HMI-Pred initially builds an ensemble classifier to screen for potential ligand sequences within microbial genomes. It then jointly evaluates sequence semantics and molecular docking to predict potential microbe-host receptor interactions.HMI-Pred achieved robust performance with F1-scores of 0.901 for microbial ligand identification and 0.883 for interaction prediction. Application to 332,381 microbial proteins revealed distinct interaction patterns: histone deacetylases (HDACs) served as broad-spectrum targets (mean score > 0.80), while G protein-coupled receptors (GPCRs) exhibited high specificity (scores 0.42–0.61). Furthermore, literature mining validated over 47% of the functional predictions, and specific immunomodulatory interactions were confirmed in Akkermansia muciniphila.HMI-Pred provides a valuable computational tool for decoding host-microbe signaling networks and facilitating the discovery of microbiome-based therapeutic targets.

**Availability:** The source code and documentation are available at https://github.com/YangLab-BUPT/HMI-Pred.

## Introduction

Bacteria play a complex, dual role in human health, functioning both as potential pathogens and as indispensable symbionts essential for maintaining host physiological functions(Chow et al., 2010; Afzaal et al., 2022). The trillions of commensal microbes colonizing the human body actively regulate immune homeostasis, defend against pathogen invasion, and promote tissue development (Curtis and Sperandio, 2011). Consequently, elucidating the molecular interactions between bacteria and host proteins represents a critical breakthrough for understanding infections, immune dysregulation, and microbiota-related diseases(Wang et al., 2021). Wang et al. demonstrated that gut bacteria express “microbial-host isozymes” that actively regulate host metabolism, revealing an unprecedented level of host-microbe enzymatic integration.(Wang et al., 2023).

Beyond metabolic enzymes, a primary mode of host-microbe interaction occurs at the cell surface, functionally constituting “cross-species intercellular communication”(Lawing and Bleich, 2025). Bacteria are capable of secreting proteins that mimic host ligands, which directly bind to host surface receptors to trigger downstream signaling(Lebeer et al., 2010). It is estimated that typical Microbe-Associated Molecular Patterns (MAMPs) involving surface receptors (such as TLRs and Nod-like receptors) account for approximately 59.2% of disease-related human-microbial protein interactions(Bhavsar et al., 2007; Zhou et al., 2022). This “crosstalk” suggests that microbes and host cells share a communication language mediated by secretory proteins and cell surface receptors.

In recent years, innovations in high-throughput technologies have revolutionized research on these cross-species interactions. Platforms such as the BASEHIT database have enabled the systematic, large-scale mapping of the relationships between humans and their commensal microbiota. By screening 3,324 human secretory proteins from four barrier tissues (gut, skin, oral cavity, and reproductive tract) against 519 bacterial strains from six major phyla, these studies have generated comprehensive databases containing thousands of interaction pairs. This provides an unprecedented resource for analyzing host–microbiota interconnections and decoding this molecular-level dialog(Sonnert et al., 2024).

Despite these experimental advances, identifying specific host-microbe interaction pairs remains a challenge due to the immense diversity of the microbiome. Traditional methods, such as 16S rRNA sequencing, primarily focus on microbial composition analysis and cannot reveal mechanistic details at the protein level(Fukuda et al., 2016; Peng et al., 2025). Therefore, computational tools capable of predicting these interactions are urgently needed. While sequence-based protein-protein interaction (PPI) prediction has made significant progress with the advent of artificial intelligence—ranging from early Multilayer Perceptron (MLP)-based models like DeepPPI to advanced architectures utilizing graph convolutional networks (e.g., S-VGAE) and self-attention mechanisms (e.g., CAMP, D-SCRIPT)—these general-purpose tools face distinct limitations when applied to the microbiome(Murakami and Mizuguchi, 2022; Du et al., 2017; Yang et al., 2020; Lei et al., 2020; Sledzieski et al., 2021). First, they often rely on sequence features while neglecting structural information (e.g., DeepFE-PPI predicts interactions but cannot identify binding interfaces, whereas TAGPPI integrates data only for proteins with known crystal structures)(Yao et al., 2019; Song et al., 2022). Second, their training data is predominantly derived from human PPIs, lacking the optimization required for host–microbiota interactions, which leads to a cross-species prediction bias.

To address these challenges and systematically decode the gut microbiota–human receptor interaction network, this study presents HMI-Pred (Host-Microbe Interaction Predictor), a novel prediction framework designed specifically for cross-species signaling communication. Unlike general PPI tools, HMI-Pred is engineered to capture the unique characteristics of microbial ligands and host receptors. The framework incorporates three primary innovations: (1) leveraging the ESM-1v protein language model to extract rich sequence features, enabling the screening of diverse ligands without reliance on crystal structures(Meier et al., 2021); (2) introducing a dual-path framework that integrates sequence semantics with biological mechanism constraints for cross-validation; and (3) utilizing the high-quality BASEHIT interaction dataset to fine-tune and optimize the prediction of bacteria–immune protein binding interfaces. By establishing an end-to-end solution from sequence to interaction probability, HMI-Pred offers a new paradigm for discovering microbiome therapeutic targets and advancing anti-infective drug development.

## Result

### Overview of HMI-Pred

In host-microbe communication, proteins on the surface of bacteria or secreted by bacteria resemble endogenous host ligands. These microbial mimics can directly bind to host cell surface receptors to modulate immune and metabolic homeostasis. Based on this biological principle, we developed HMI Pred—an end-to-end computational pipeline. This pipeline first identifies microbial proteins that resemble host ligands, and then systematically evaluates the interaction potential between these protein ligands and human receptors (shown in Figure 1). The HMI-Pred framework consists of three core modules.

**Fig 1.**
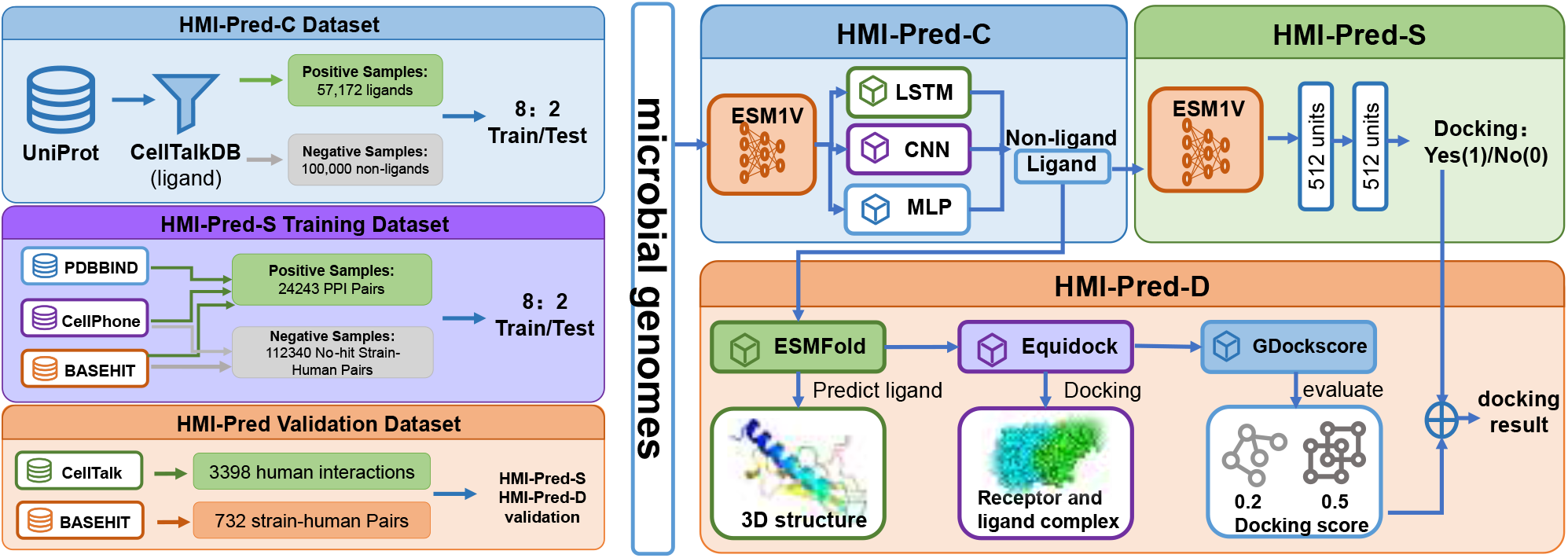
Overview of HMI-Pred’s dataset and pipeline.HMI-Pred-C is used to identify whether sequences belong to human ligands, while HMI-Pred-S and HMI-Pred-D assess the likelihood of microbe-host protein interactions from sequence-level and molecular docking mechanism perspectives, respectively.

The HMI-Pred-C (Classification) module screened ligands that similar to human ligands from microbial genome. HMI-Pred-C integrated three neural network architectures (Taud and Mas, 2017; Chua and Roska, 2002; Greff et al., 2016), with fixed weights of 0.4 for the MLP, 0.3 for the CNN, and 0.3 for the LSTM. The HMI-Pred-D (Protein-Protein Docking) Module predicted the 3D structures of ligands and receptors using ESMFold, followed by protein-protein docking using EquiDock(Lin et al., 2023; Ganea et al., 2021). The quality of docking conformations was evaluated using GDockScore(McFee and Kim, 2023). The HMI-Pred-S (Sequence-based Scoring) Module extracted sequence features of the ligands and receptors using protein large language model ESM-1v to construct 1280-dimensional feature vectors. The model outputs from HMI-Pred-D and HMI-Pred-S were integrated using a customized scoring function to comprehensively assess protein ligand-receptor interconnections.

### Performance of different models in gut microbial ligand protein identification

To validate the model’s capability in identifying authentic signaling molecules, we employed the CellTalk dataset (Shao et al., 2021), a comprehensive repository of curated human ligand–receptor interactions. The evaluation results on this test set demonstrated that the DL-based integrated learning strategy of HMI-Pred-C exhibited marked advantages over traditional machine learning (ML) algorithms in identifying potential human ligand sequences (Table 1). Among the conventional ML methods, SVM achieved maximum performance with precision, recall, and F1-scores of 0.743, 0.807, and 0.774, respectively, followed by XGBoost and SGD(Hearst et al., 1998; Chen, 2016; Amari, 1993). The Decision Tree algorithm showed a relatively lower performance across all metrics (0.672, 0.686, and 0.679). The DL(deep learning) models substantially outperformed traditional ML approaches. The MLPs demonstrated excellent performance across all three metrics (0.863, 0.937, and 0.899). CNN and LSTM networks delivered similar outstanding results, with F1-scores reaching 0.884 and 0.890, respectively. The ensemble model combining MLP, CNN, and LSTM exhibited a superior comprehensive performance: while maintaining high precision (0.842), it achieved exceptional recall (0.970), resulting in an optimal F1-score (0.901).

**Table 1.** Performance of different models in ligand identification.

| Model | Precision | Recall | F1-Score |
| --- | --- | --- | --- |
| SVM | 0.743 | 0.807 | 0.774 |
| Decision Tree | 0.672 | 0.686 | 0.679 |
| SGD | 0.688 | 0.757 | 0.721 |
| XGBoost | 0.694 | 0.792 | 0.740 |
| MLP (t=0.5) | 0.863 | 0.937 | 0.899 |
| CNN | 0.870 | 0.899 | 0.884 |
| LSTM | **0.878** | 0.903 | 0.890 |
| Ensemble model<br>(MLP+CNN+LSTM) | 0.842 | **0.970** | **0.901** |

### Evaluation of gut microbe-host receptor interactions prediction

To assess the efficacy of utilizing one-dimensional sequence information alone, the performance of the sequence-based scoring module (HMI-Pred-S) was evaluated against two baseline models: a Random Forest (RF) model utilizing ESM1v-extracted features and a neural network model employing one-hot encoding. As illustrated in Figure 2, the HMI-Pred-S model demonstrated superior predictive capability in identifying ligand–receptor pairs from external datasets (sourced from CellPhoneDB and CellTalkDB, which were excluded from training) (Efremova et al., 2020; Shao et al., 2021). At a threshold of 0.5, the neural network-based model achieved high accuracies of 0.955 and 0.724, respectively. In sharp contrast, when compared to the Random Forest model (Breiman, 2001), the performance gap was evident, as the traditional algorithm yielded significantly lower accuracies of 0.355 and 0.514.

**Fig 2.**
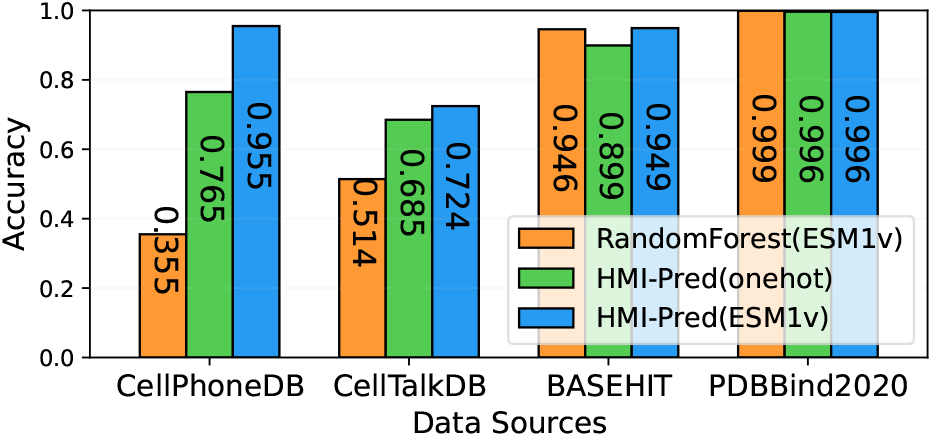
Performance of HMI-Pred under different feature encodings and models

Subsequently, the contribution of three-dimensional structural information was evaluated to determine if integrating docking features could further enhance predictive performance. An ablation study was conducted comparing three configurations: the docking-based module (HMI-Pred-D), the sequence-based module (HMI-Pred-S), and their integrated framework (HMI-Pred-S&D). These models were assessed using the CellTalkDB and the experimentally verified BASEHIT datasets, alongside overall precision, recall, and F1 scores on combined test sets.

Comparative analysis indicated that the integrated HMI-Pred-S&D model consistently outperformed the individual modules across all metrics (Table 2). Notably, the recall rate improved remarkably to 0.847 in the combined model, significantly surpassing both HMI-Pred-D (0.552) and HMI-Pred-S (0.632). Ultimately, the integrated framework achieved an F1 score of 0.883. Furthermore, when benchmarked against FoldDock—a baseline tool incorporating AlphaFold for prediction and docking—the proposed HMI-Pred framework demonstrated significant advantages. As indicated in Table 2, HMI-Pred achieved improvements in various metrics exceeding 15%, particularly in the accurate identification of experimentally verified interactions within the BASEHIT dataset (0.822 vs. 0.547).

**Table 2.** Performance comparison of different prediction methods on protein interaction tasks.

| Metrics | Prediction Methods |  |  |  |
| --- | --- | --- | --- | --- |
|  | HMI-Pred-D | HMI-Pred-S | HMI-Pred-S&D | FoldDock |
| CellTalk accuracy | 0.4946 | 0.7243 | **0.8493** | 0.5059 |
| BASEHIT accuracy | 0.7587 | 0.6570 | **0.8222** | 0.5471 |
| Test Precision | 0.848 | 0.886 | **0.925** | 0.778 |
| Test Recall | 0.552 | 0.632 | **0.847** | 0.520 |
| Test F1 | 0.684 | 0.743 | **0.883** | 0.634 |

### Large-Scale Analysis of Gut Microbiota–Human Receptor Interactions

Based on this computational pipeline, the interactions between human gut microbiota and host receptor proteins were systematically evaluated. This study downloaded 332,381 gut microbe-related protein sequences from the NCBI database(https://www.ncbi.nlm.nih.gov/), and initially screened 35,683 potential ligand proteins using the HMI-Pred-C model. Subsequently, the HMI-Pred-D and HMI-Pred-S models were employed to systematically analyze the interactions between these ligand proteins and key human receptors, including epigenetic regulatory receptors (HDAC1-9), and G protein-coupled receptors (GPR41/GPR43/TGR5/GIPR)(de Ruijter et al., 2003; Rosenbaum et al., 2009). These play crucial roles in physiological processes such as immune regulation and metabolic control. PPI scores were calculated to identify gut bacterial strains capable of specific interactions with these receptors.

To systematically characterize the potential of gut microbiota in modulating human signaling pathways, the interaction landscape was visualized by specifically selecting the bacterial species that exhibited the highest predicted interaction score for each human receptor. (Figure 3A). Hierarchical clustering revealed a distinct high-affinity receptor cluster predominantly composed of histone deacetylases (HDACs), specifically HDAC1, HDAC3, HDAC6, HDAC7, and HDAC8. These receptors exhibited consistently high interaction scores across diverse bacterial phyla (mean score of the top 20 species > 0.80), suggesting that the modulation of host epigenetic machinery is a broad-spectrum and evolutionarily conserved trait among these dominant gut commensals. In contrast, G-protein-coupled receptors (GPCRs), such as TGR5 and GPR41, clustered into a low-affinity group characterized by significantly lower interaction scores (0.42–0.61), indicating a higher degree of ligand specificity or a more restricted range of bacterial activators.

**Fig 3.**
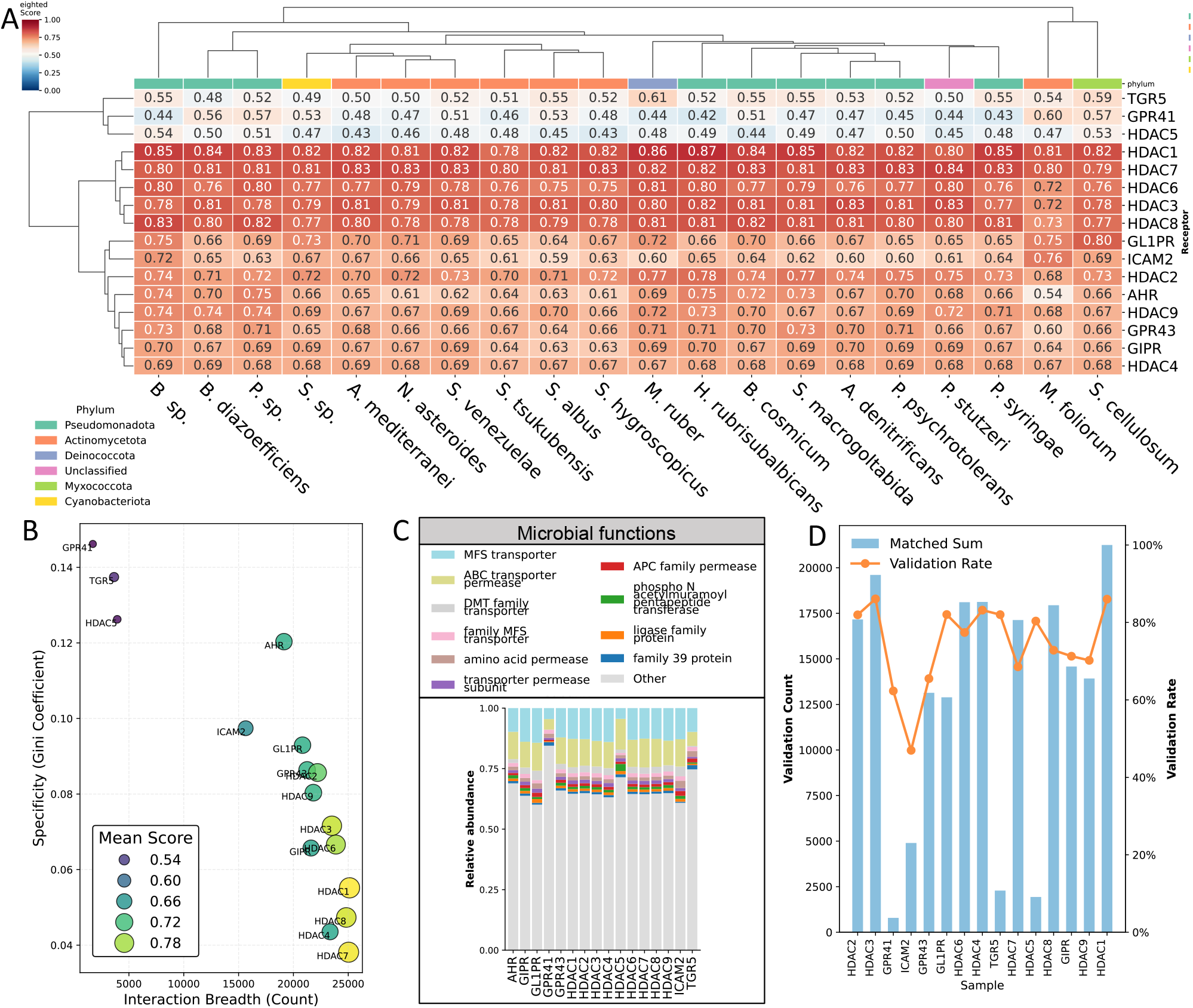
Comprehensive analysis of gut microbiota-host receptor interactions. A.Heatmap of the highest predicted interaction scores between bacterial species and human receptors. B. Relationship between interaction breadth and receptor specificity. C. Functional enrichment of bacterial strains associated with receptor binding. D. validation rate of differentreceptors.

Taxonomically, high-scoring species were predominantly enriched within the phyla Actinomycetota and Pseudomonadota. Notable functional redundancy was observed across phylogenetically distant lineages; for instance, Streptomyces species (e.g., S. hygroscopicus, S. albus) from Actinomycetota and Pseudomonas species (e.g., P. syringae, P. stutzeri) from Pseudomonadota both displayed potent interaction profiles with the HDAC cluster. Furthermore, specific species such as M. ruber and H. rubrisubalbicans emerged as “interaction hubs,” demonstrating robust binding potential in multiple receptor families, including HDACs and specific GPCRs such as GL1PR and ICAM2.

To quantify the selectivity of host-microbe protein interaction, Figure 3B illustrates the relationship between interaction breadth (defined as the count of high-scoring bacterial species) and receptor specificity (measured by the Gini coefficient)(Dorfman, 1979). The receptor proteins segregate primarily into two distinct categories. The first, identified as a “broad-spectrum” cluster, is predominantly composed of histone deacetylases (HDACs) and specific metabolic receptors. Receptors such as HDAC1, HDAC7, and HDAC8 occupy the lower-right quadrant, characterized by extensive interaction breadth (> 20, 000 species) and minimal Gini coefficients (< 0.06). In contrast, receptors like GPR41 and TGR5 exhibit a “high-specificity” profile, demonstrating restricted interaction spectra (breadth < 5, 000) and elevated Gini coefficients (> 0.12). Their lower average interaction scores suggest that their activation is highly selective, likely contingent upon specific ligands produced by a limited subset of bacterial taxa rather than ubiquitous metabolites. Transitional receptors, such as AHR and ICAM2, occupy an intermediate region, bridging the gap between highly specific GPCR signaling and broad-spectrum epigenetic modulation. Figure 3C presents analysis of the functional characteristics of these bacterial strains, demonstrating that gut microbes with transporter and permease functions are more likely to interact with receptor proteins.

Further analysis revealed that gut microbial proteins with transporter, permease, transferase, and ligase functions were particularly strongly associated with human immune-related proteins in our dataset. To systematically validate these predictions, we retrieved PubMed(http://www.pubmed.org) literature associated with each predicted ligand protein and analyzed their abstracts for functional annotations. A prediction was considered validated if the abstract explicitly mentioned the function of the corresponding microbial protein. This conservative approach yielded an overall verification rate > 47% across all receptor types (Figure 3D), indicating that nearly half of our functional predictions are supported by experimental evidence. The remaining may either represent novel interactions not yet reported in the literature or FPs obtained from our computational pipeline. Notably, the verification rate varied among different receptor families, with HDAC-interacting proteins being particularly robustly supported by publications. These validation results confirm that our integrative computational approach can reliably identify biologically relevant gut microbe–host protein interactions while also highlighting potentially novel interactions worthy of experimental investigation.

### Validation of Interactions Between Akkermansia muciniphila and Human Immune Proteins via HMI-Pred

The mucin-degrading bacterium, *Akkermansia muciniphila*, has garnered significant attention due to its beneficial roles in host metabolism, immune regulation, and gut barrier function(Zheng et al., 2023). To validate the predictive power of our HMI-Pred platform, we focused on two well-characterized immunomodulatory proteins from this bacterium: Amuc 1100 and Amuc 2109. These proteins interact with the host receptors GPR41, GPR43, and TLR2, promoting regulatory T cell differentiation and modulating NF-kB signaling pathways.

Our validation approach began with the retrieval of complete *A. muciniphila* genomes from NCBI, followed by a systematic screening with HMI-Pred-C to identify potential ligand proteins. Sequence alignment against the reference proteins Amuc 1100 and Amuc 2109, utilizing DIAMOND with a 20% identity threshold, yielded 49 homologs, suggesting that these putative ligands may share functional properties with known immunomodulators. Remarkably, HMI-Pred-based analysis predicted that all 49 candidate sequences could interact with ≥ 1 of the target receptors (TLR2, GPR43, or GPR41), with 27 sequences showing binding potential with ≥ 2 receptor types.

## Method

### HMI-Pred pipeline

This study proposed a novel, ligand–receptor interconnection prediction pipeline named HMI-Pred to systematically evaluate the interactive potential between protein ligands and human receptor proteins. Unlike traditional ligand–receptor-based docking methods, this model innovatively incorporated a bacterial strain–human interrelationship discrimination module, enabling a more comprehensive assessment of the binding probability between ligands and human proteins. The HMI-Pred framework consists of three core modules, discussed below.

To accurately screen potential human-protein-related ligands, the HMI-Pred-C module employs a deep ensemble learning framework that integrates three distinct neural architectures via a dynamic weighting strategy. Specifically, the framework comprises a three-layered Multilayer Perceptron (MLP) with 256-64-1 units, ReLU activation, and a dropout rate of 0.5; a 1D Convolutional Neural Network (1D CNN) consisting of three convolutional blocks (kernel sizes 7, 5, and 3) followed by batch normalization, max-pooling, and a 128-1 fully connected output; and a bidirectional bi-layered Long Short-Term Memory (LSTM) network with a hidden dimension of 128 and a dropout rate of 0.3 to capture sequential dependencies. To effectively fuse these models, we introduced learnable weight parameters initialized uniformly and normalized via the Softmax function, which dynamically adjusted the ensemble distribution during training to maximize numerical stability, ultimately converging to fixed weights of 0.4 (MLP), 0.3 (CNN), and 0.3 (LSTM). To benchmark the efficacy of this proposed framework, we conducted comparative experiments against five traditional machine learning algorithms: Extreme Gradient Boosting (XGBoost), Random Forest (RF), Decision Tree (DT), Support Vector Machine (SVM), and Stochastic Gradient Descent (SGD). For fair comparison, hyperparameters for tree-based models were standardized (n estimators=300, max depth=8), and learning rates for XGBoost and SGD were fixed at 0.1. All models were rigorously evaluated using 5-fold cross-validation to ensure robust generalization capability.

Following the preliminary screening, the framework employs a dual-branch strategy to comprehensively evaluate the interaction capacity between ligand and receptor. The structural branch, HMI-Pred-D, assesses physical interaction potential by first predicting the 3D conformations of ligands and receptors via ESMFold, followed by rigid-body docking using EquiDock. The quality of the resulting docking conformations are rigorously quantified using the GDockScore. HMI-Pred-S, the sequence-based branch, captures semantic interaction features utilizing the ESM-1v model to extract 1280-dimensional feature vectors from ligand and receptor sequences, which are subsequently processed by a bi-layered neural network (512 and 64 hidden units) to estimate interaction probabilities. To yield a holistic prediction, the structural affinity metrics from HMI-Pred-D and the sequence-based probability scores from HMI-Pred-S are fused through a customized scoring function, thereby ensuring a robust assessment of protein ligand-receptor interconnections.

### Datasets

To accurately identify the human-protein-related ligands, based on the ligand protein catalog provided by the CellTalkDB database, the positive ligand samples were obtained from the reviewed entries available in UniProt, yielding 57,172 positive samples with high confidence. For the negative samples, reviewed non-ligand proteins, 100,000 in number, were randomly selected. The training and testing sets at an 8:2 ratio.

For preparing training data for the HMI-Pred-S Module, a multi-source biomolecular interaction dataset was constructed for sequence-based interlinkage prediction. The positive samples were integrated from three sources: 19,037 protein-protein interaction (PPI) pairs from the PDBBind database, 1,877 experimentally validated ligand–receptor pairs from the CellPhoneDB database, and 3,730 strain–human protein interaction pairs (classified as strong- or weak-hits) from the BASEHIT database(wwPDB consortium, 2019). The negative samples were composed of 62,340 non-interacting (no-hit) strain–human protein pairs extracted from the BASEHIT database. Additionally, to augment the negative dataset, 50,000 non-redundant random protein pairs were generated by shuffling proteins sourced from CellPhoneDB. This composite dataset was subsequently partitioned into training and testing subsets at an 8:2 ratio.

For the Evaluation Dataset for HMI-Pred-D and Overall HMI-Pred Validation, the performance of the PPD pipeline was assessed. It used the 3398 ligand–receptor complexes from the CellTalkDB database (evaluating human protein interconnections), and 732 test strain–human protein interaction pairs from the BASEHIT database (assessing microbial protein binding to human receptors). These test sets were employed to validate the accuracy of structure-based docking and integrated interaction prediction workflows.

### Evaluation metrics

The performance of the HMI-Pred framework was systematically evaluated with a comprehensive set of metrics across all modules. For the HMI-Pred-C ligand identification module, three key evaluation metrics were employed: Precision, which reflected the reliability of positive predictions; Recall, measuring the model’s ability to identify truly positive ligands; and the F1-score, representing the harmonic mean of precision and recall. In this evaluation, True Positives (TPs) were defined as correctly identified ligand proteins, False Positives (FPs) as non-ligand proteins incorrectly classified as ligands, and False Negatives (FNs) as actual ligands missed by the model. To thoroughly assess model performance, the integrated learning approach and baseline methods were compared through multiple metrics.

The evaluation of the HMI-Pred-S sequence-based scoring module primarily utilized accuracy as the key performance-determining metric, calculated as the ratio of correct (TP + TN) to total predictions (TP + FP + TN + FN). To rigorously test the generalization capability of the model across various biological contexts, four distinct datasets were independently validated: the ligand protein collection from CellTalkDB, strain–human protein interactions from BASEHIT, experimentally verified ligand–receptor pairs from CellPhoneDB, and PPIs from PDBBind2020. The variations in model performance across these diverse datasets provided critical insights into its robustness for predicting PPIs under various scenarios.

A series of carefully designed ablation experiments validated the overall effectiveness of the complete HMI-Pred pipeline. The independent contributions of the sequence feature analysis (HMI-Pred-S) and structural docking (HMI-Pred-D) pathways using the CellTalkDB and BASEHIT datasets was quantified, respectively. During the benchmark testing phase, the model’s baseline performance was established by calculating the overall accuracy metrics for these two datasets. For a more detailed evaluation, a combined positive sample set was created by merging all CellTalkDB samples with strong- and weak-hit interactions, while the negative sample set consisted exclusively of no-hit pairs from BASEHIT. This comprehensive evaluation framework employed precision, recall, and F1-score metrics to thoroughly assess model performance. A comparison between the full model and its simplified variants effectively demonstrated the essential contribution of each module and the synergistic impacts of all modules within the complete pipeline.

### Analysis of Interactions Between Gut Microbial Genomes and Human Receptor

HMI-Pred was employed to evaluate the interaction potential of 332,381 microbial protein sequences retrieved from the NCBI database with key human receptors. To rank the regulatory capacity of specific bacterial taxa, a composite score was computed to integrate both interaction strength and ligand abundance:

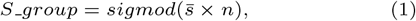

where 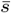 denotes the average interaction score and *n* represents the count of ligand proteins within the group. Receptor specificity was quantified utilizing Gini coefficients and hierarchical clustering. Functional annotations (e.g., transporters) were analyzed, and biological relevance was systematically validated by mining PubMed abstracts for explicit functional corroboration of the predicted microbial proteins.

## Discussion

In this study, HMI-Pred is presented as a novel end-to-end deep learning framework designed to systematically decode the interactions between the gut microbiota and the host. By innovatively integrating the protein language model ESM-1v with structural docking mechanisms (ESMFold and EquiDock), the limitations associated with the excessive reliance of traditional methods on sequence similarity or the prohibitive computational costs of crystal structure simulations are effectively overcome. The model was systematically evaluated on the CellTalkDB and CellPhone datasets, as well as on the experimentally verified BASEHIT interaction dataset. Comprehensive results indicate that the unique characteristics of microbial ligands and host receptors are successfully captured by HMI-Pred, thereby establishing a robust paradigm for the identification of microbe-host interactions in the absence of a prior structural knowledge.

The superior performance of HMI-Pred is rendered particularly evident when benchmarked against existing structural docking baseline models. As illustrated in Table 2, the integrated HMI-Pred-S&D model is shown to significantly outperform FoldDock across multiple evaluation metrics. Specifically, on the CellTalkDB, an accuracy of 0.849 was achieved by HMI-Pred, which is markedly higher than the 0.506 recorded by FoldDock; meanwhile, on the more challenging, real-world host-microbe interaction dataset (BASEHIT), a substantial advantage was maintained (0.822 versus 0.547). This performance disparity suggests that, while AlphaFold-based methods (such as FoldDock) are powerful in precise structural determination, the semantic adaptability required for identifying host-microbe binding interfaces may be lacking when relying solely on structural perspectives. A more error-tolerant and biologically relevant predictive capability is provided by the dual-path strategy adopted in HMI-Pred—wherein sequence semantics are fused with physical docking constraints—effectively mitigating the cross-species prediction bias that is frequently observed in models trained exclusively on human PPI data. Furthermore, based on the analysis of the interaction landscape depicted in Figure 3A, it is revealed that a robust binding potential with critical host regulatory proteins is exhibited by specific bacterial taxa—a finding that further substantiates the biological validity of the high-scoring predictions. For instance, high-affinity interactions between histone deacetylases (HDACs) and the *Pseudomonas* and *Streptomyces* genera (exemplified by species such as *P. stutzeri* and *S. hygroscopicus*) were successfully identified by the model. Literature validation supports that these interactions are mediated by direct surface-level protein engagement rather than solely by soluble metabolites. For instance, species within the *Pseudomonas* genus (e.g., *P. stutzeri*) are known to express surface pili and outer membrane proteins (OMPs) that act as adhesins. These surface proteins directly bind to host cell receptors to facilitate colonization and cellular signaling(Uszczynski et al., 2019). With respect to pathogens, *N. asteroides* is verified as a human pathogen capable of invading through multiple routes and causing suppurative inflammation and abscess formation in various tissues(Fatahi-Bafghi, 2018). These examples demonstrate that our computational approach successfully identified both beneficial and pathogenic bacteria with documented human interactions, thereby supporting the biological relevance of our screening strategy.

To exlpore the interactions between probiotics and human receptors, the interaction profiles of *Lactobacillus* and *Bifidobacterium* were analyzed(shown in Figure 4). Distinct functional differentiation was revealed by the results: preferential affinity for immune and barrier-related receptors (e.g., ICAM2, AHR) was exhibited by *Bifidobacterium*, consistent with its established role in enhancing mucosal integrity(Abdulqadir et al., 2023; Barreira-Silva et al., 2025); whereas a stronger binding potential for metabolic receptors (e.g., GL1PR, GIPR) was demonstrated by *Lactobacillus*, implying a direct involvement in the regulation of host glucose homeostasis(Ke et al., 2025; Wang et al., 2024). Notably, robust interactions with HDACs were maintained by both genera, supporting a shared pathway of broad epigenetic modulation mediated by microbial metabolites. it can also be observed that *L. acidophilus* can interact with most receptors, while *B. bifidum* more easily interacts with AHR and HDAC proteins. This pattern highlights that the health benefits of probiotics are likely driven collectively by broad epigenetic regulation and highly specific receptor-mediated immune modulation.

**Fig 4.**
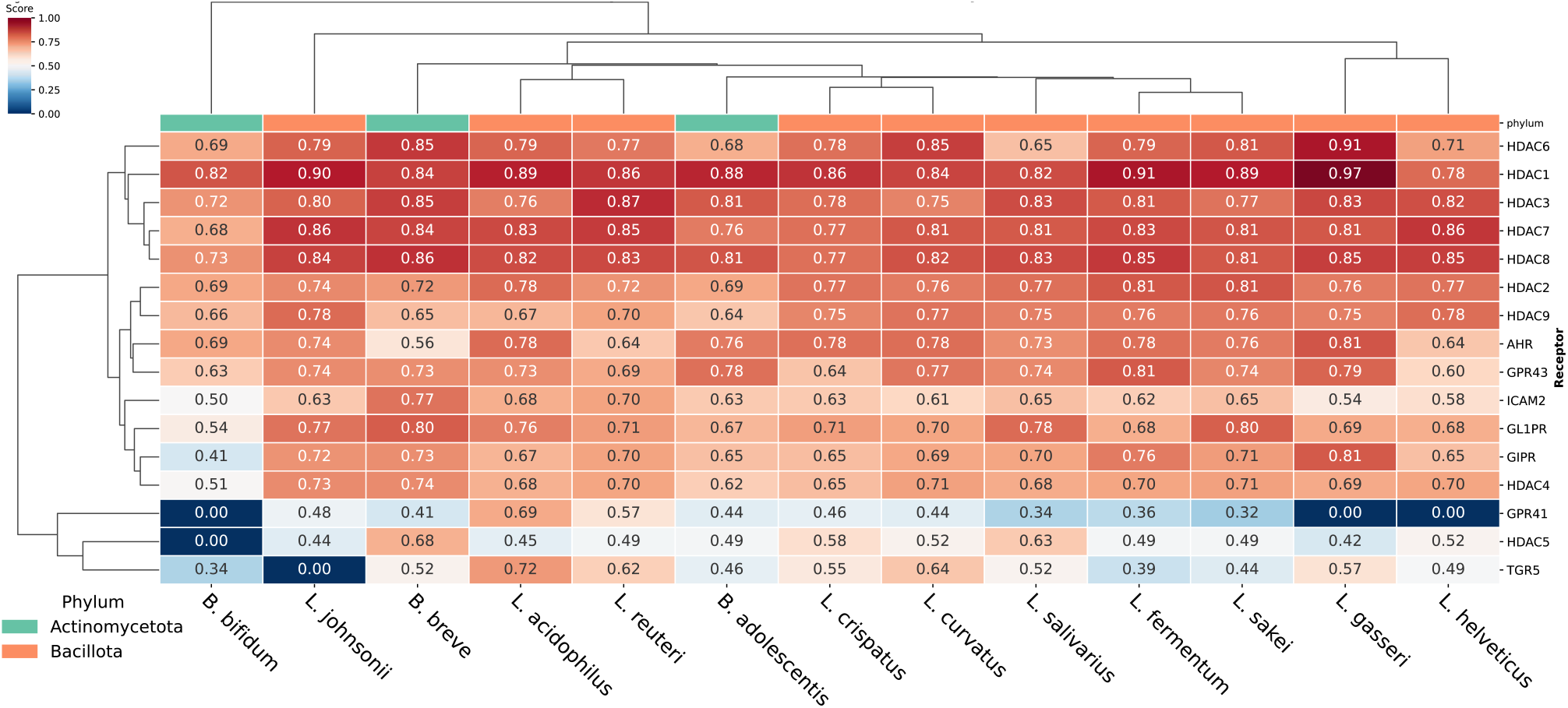
Interaction scores between Lactobacillus/Bifidobacterium and host receptors.

The model is currently constrained by the limited scale of experimentally verified data and its exclusive focus on protein-protein interactions, neglecting the critical regulatory role of microbial metabolites. Future work will aim to address these limitations by developing a multi-modal framework that integrates metabolomics data and expands training resources to achieve a more comprehensive decoding of host-microbe communication.

## Conclusion

This study presents HMI-Pred, a specialized deep learning framework for predicting potential interactions between gut microbiota and human receptors, by integrating sequence semantic information with ligand-receptor structural docking. Our approach’s performance was verified in public cell communication datasets, and real microbe-host protein interaction data. HMI-Pred can be a valuable computational tool for analyzing host-microbe protein interactions and help microbiologists to decode gut microbe-host molecular-level dialogue.

## Author contributions statement

HL drafted the work and contributed to the creation of new software used in the work. RZ and CM contributed to the interpretation of data.TC and RJ contributed to the acquisition. YY and XL substantively revised the draft. All authors have approved the submitted version and have agreed both to be personally accountable for the author’s own contributions. All authors read and approved the final manuscript.

## Acknowledgments

The authors thank the anonymous reviewers for their valuable suggestions. This work is supported in part by funds from the National Natural Science Foundation of China (NSF: 62203060 and 62403492).

